# CH···O Interactions Are Not the Cause of Trends in Reactivity and Secondary Kinetic Isotope Effects for Enzymatic S_N_2 Methyl Transfer Reactions

**DOI:** 10.1101/071043

**Authors:** Jianyu Zhang, Judith P. Klinman

## Abstract

**Compaction matters:** S_N_2 substitution represents an important class of reaction for both chemical and biological systems. The ability to assess enzymatic transition state structure within this class of reaction remains a major experimental challenge. Here, we comment on and compare the relative impact of compaction along the axis of reaction to the impact of an orthogonal CH···O hydrogen bonding interaction. The latter is concluded to play a limited role in determining relative reaction rates and secondary KIEs derived from experimental structure-activity correlations.

---

Understanding and predicting chemical reactivity remains one of the central tasks of molecular science. Much attention has been focused on the properties of reactants and, in particular, the concept of “structure-reactivity relationships” whereby changes in structure lead to changes in reactivity. Extension of such relationships, from either computational or experimental gas phase measurements to the condensed phase, introduces important issues regarding the impact of environment on reactivity. This becomes especially important for enzymes where evolved protein structures produce incredibly accelerated reaction rates. In a recent work, Williams and coworkers carried out DFT calculations on a simplified model reaction to understand the influence of carbon to oxygen hydrogen bonds (CH···O) on the magnitude of rates and kinetic isotope effects (KIE) in enzymatic methyl transfer reactions.^[1]^ They suggested that CH···O interactions could play a dominant role in these effects. However, as shown below, the role of such hydrogen bonding is concluded to be minor, while compactness/tightness along the axis of the reaction (donor-acceptor distance) is predicted to have the major impact on the magnitude and trend of secondary KIEs.

For a typical S_N_2 reaction, the event of bond breaking and bond forming occurs synchronously, forming a pseudo penta-coordinate transition state (TS) in which both nucleophile and leaving group are partially bonded to the central transferred group. For methyl transfer, the structure of the TS at the α sp^3^ reacting carbon center can be symmetric, early (reactant like) and late (product like). Determination of this TS structure is key to elucidating the reaction mechanism and has received an enormous amount of attention ^[2][3]^. In particular, primary carbon and secondary α-deuterium/tritium KIEs (2° α-D/T) have played a dominant role in mechanistic reconciliations.

As described by Trievel and coworkers,^[4]^ CH···O hydrogen bonding interactions between active site side chains and the activated methyl group of AdoMet are predicted to influence reaction outcomes within the family of AdoMet -dependent methyltransferases. The precise catalytic role of such CH···O hydrogen bonding on TS structure and the size of KIE has been recently analyzed by Williams and coworkers^[1]^ using DFT simulations to interrogate the enzymatic system catechol O-methyltransferase, COMT. Their (simplified) methyl transfer model contains a methyl cation surrounded by two axial water molecules and three equatorial water molecules; these water molecules mimic the donor-acceptor interaction along the axis of reaction (referred to as axial interactions) and the hydrogen bonding interaction between the transferred CH_3_ and hydrogen bonding functional groups in an orthogonal plane (referred to as equatorial interactions). By first fixing the axial donor-acceptor distance at 4 Å (r_ax_ =2 Å), the impact of the equatorial CH···O interaction on the simulated 2° α-D_3_ KIE was found to be >1 in all cases; importantly, the magnitude of this KIE became more normal as the extent of H-bonding increased: KIE = 1.160 (when r_eq_ =3 Å) vs. KIE = 1.121 (when r_eq_ =4 Å) corresponding to a KIE change of −0.039/ Å , Table 1. At the same time, the energy of activation for reaction was predicted to decrease from 23.2 kJ/mol (r_eq_ = 4 Å) to 17.5 kJ/mol (r_eq_ = 3 Å), with the implication that increased hydrogen bonding would lead to increased catalytic rate. One feature of this analysis is the exclusive focus on stretching frequencies, with a lack of analysis of the out-of-plane bending vibrations; in general it has been concluded that changes in out-of-plane bending vibrations will determine the trend in 2° α-D KIE for S_N_2 reaction, with changes in vibrational frequencies deciding the magnitude of the KIE ^[2a,2d]^.

**Table 1.**
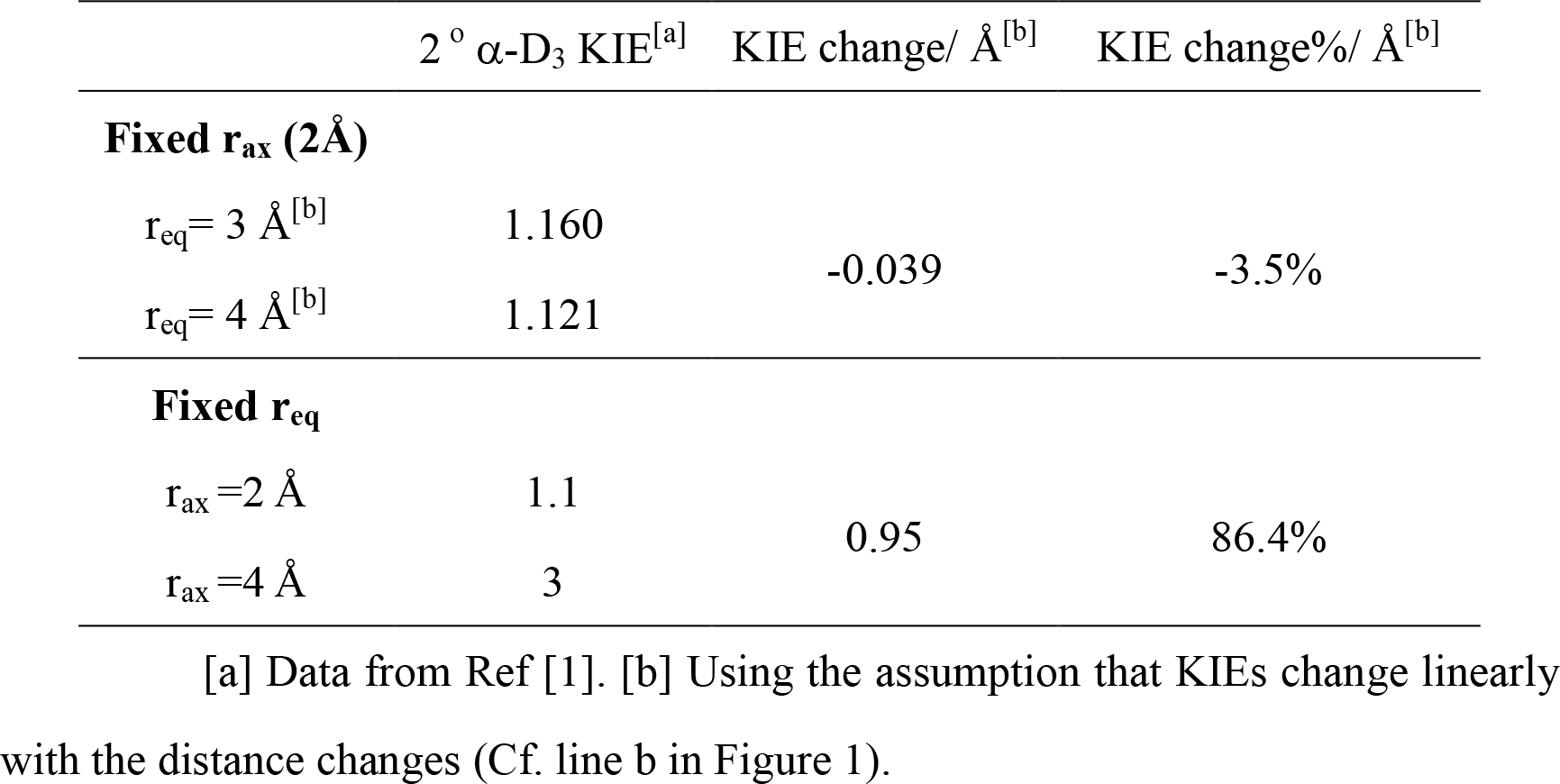
Secondary α-D_3_ kinetic isotope effects (2° α-D_3_ KIE) and their sensitivity to changes of donor-acceptor distance (2×r_ax_) and CH···O bond distance (r_eq_) for simulations of methyl transfer within a water-constrained cage.

Using the same simulation method, the magnitude of the 2° α-D_3_ KIE without any CH···O hydrogen bonding interactions has been calculated to be 1.1 and 3 when the r_ax_ is equal to 2 Å and 4 Å, respectively, showing a KIE change of 0.95/Å. Compared to the change of KIE along the equatorial coordinate (- 3.5%/Å), the change is more than 20 time larger along the axial coordinate (86.4%/Å). Under the assigned boundary conditions used to simulate the behavior of the native COMT, it is quite clear that the secondary KIE will be much more sensitive to any distance change between the methyl donor and acceptor than to changes in CH···O hydrogen bonding interaction, i.e., the major contributor to the magnitude of the KIE will come from the donor-acceptor distance (2 × r_ax_) that reflects the compactness/tightness at the reaction center.

As computed by Williams and co-workers, a decrease in either the CH···O hydrogen bonding interaction (corresponding to an increase in r_eq_) or the donor-acceptor interaction (increase in r_ax_) leads to an increase the barrier height (Table 2). At each fixed value for r_ax_, an increase in r_eq_ by 1 Å leads to a barrier height ratio around 1.2. This value is much smaller than the ratio of barrier heights at fixed r_eq_ for an increase in r_ax_ by 1 Å (ratio around 5.6). As with the magnitude of the secondary KIEs, the trends in activation energy reveal that the donor-acceptor distance along the reaction axis, rather than the equatorial CH···O hydrogen bonding interaction, will play the major role in determining catalytic rate enhancement.

**Table 2.** Barrier height (unit kJ mol^−1^) ^[a]^ for different donor-acceptor distances (r_ax_) and CH···O bond distances (r_eq_).

| $r_{eq}$<br>$r_{ax}$ | 3.0 Å | 3.5 Å | 4.0 Å | Ratio of barrier height<sup>[b]</sup> |
| --- | --- | --- | --- | --- |
| 2.0 Å | 17.5 | 20.2 | 23.2 | *1.3* |
| 2.5 Å | 66.6 | 70.9 | 79.4 | *1.2* |
| 3.0 Å | 104 | 113 | 128 | *1.2* |
| Ratio of barrier height<sup>[c]</sup> | *5.9* | *5.6* | *5.5* |  |

[a] Data from Ref [1]. [b] For $r_{eq}$ 4 Å / $r_{eq}$ 3 Å as a function of $r_{ax}$ , numbers in *italics* [c] For $r_{ax}$ 3 Å / $r_{ax}$ 2 Å as a function of $r_{eq}$ , numbers in *italics*.

Finally, a particularly compelling mechanistic insight regarding the dominance of axial vs. equatorial interactions comes the experimental observations of an approximately linear correlation between the experimental rate constant for substrate methylation and the magnitude of the observed 2° KIE within a series of site specific mutants of COMT^[5]^. In these experimental studies, there is a regular increase in the magnitude of the α 2° KIE with a decrease in rate (negative slope), which is opposite to the predictions of Williams et al. in which the KIE is expected to become less normal as the CH· · ·O elongates and the rate goes down.

To conclude, the recent study of Williams et al. has illustrated the powerful potential of computation to assess a proposed role for C-H to oxygen interactions in modulating rate and TS structure for enzymatic methyl transfer reactions. While this specific interaction has been shown to be capable of producing significant rate accelerations, its impact appears quite small relative to axial, donor-acceptor distance effects.^[1]^ Further, the experimental trends in reaction rates and KIEs[^5^] are opposite to those predicted for the equatorial H-bonding effects and in strong support of a dominance of axial donor-acceptor distances as a major correlate of enzymatic efficiency.

**Figure 1.**
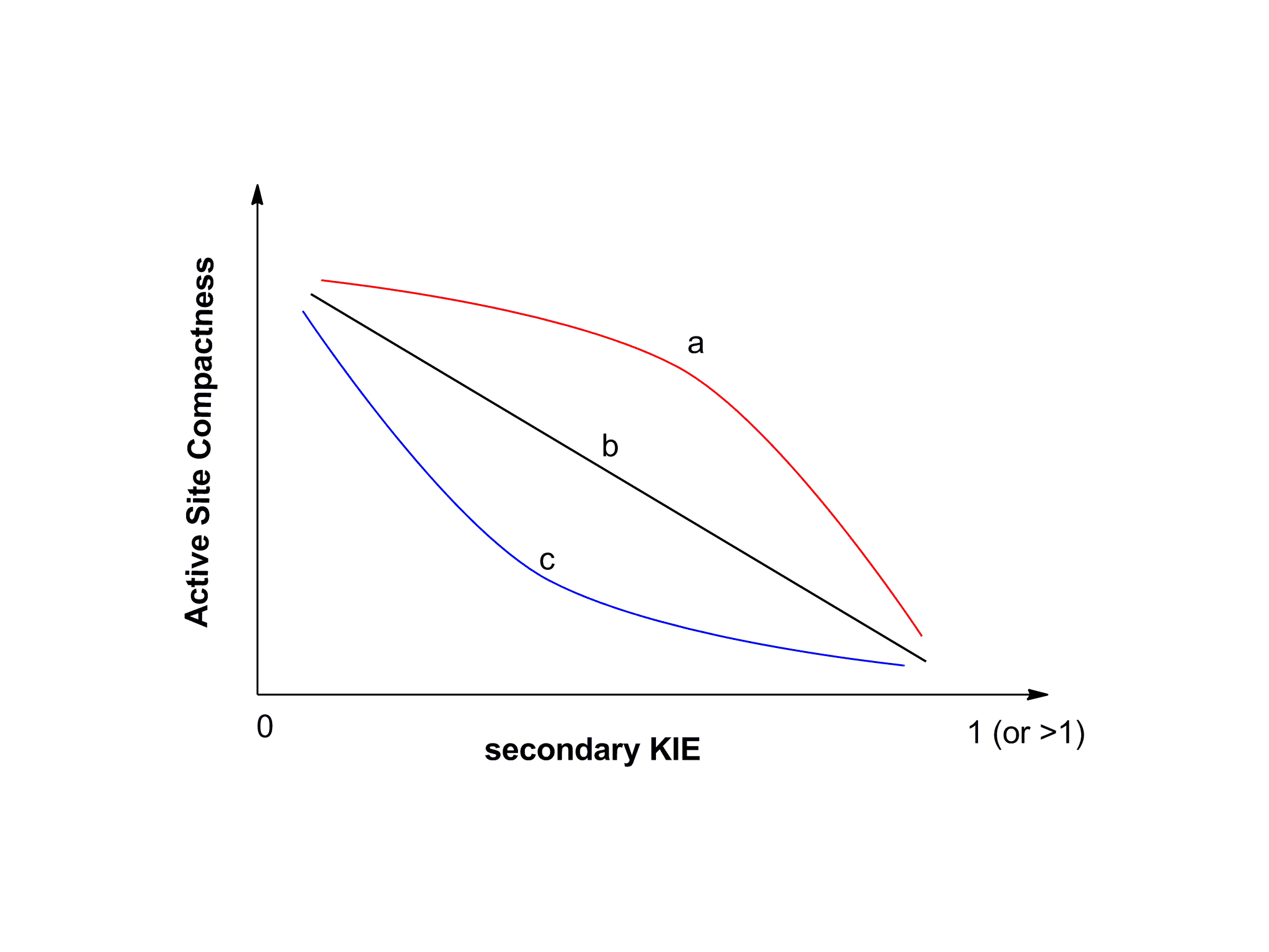
The relationship between the compactness of reaction center and the secondary KIE is currently at an empirical stage. It is expected that rigorous mathematical analysis will be forthcoming in the future developments.

